# Molecular Mechanisms Governing Peptide Nanodisc Assembly and Stability

**DOI:** 10.64898/2026.03.18.712699

**Authors:** Bikash R. Sahoo, Bankala Krishnarjuna, Thirupathi Ravula, G. M. Anantharamaiah, Ayyalusamy Ramamoorthy

**Affiliations:** Biophysics Program, The University of Michigan, Ann Arbor, MI 48109, United States; Department of Chemistry, The University of Michigan, Ann Arbor, Michigan 48109, United States; Biomedical Engineering, Michigan Neuroscience, Macromolecular Science and Engineering, The University of Michigan, Ann Arbor, MI 48109, United States; Department of Medicine, University of Alabama at Birmingham Medical Center, Birmingham, AL, 35294, United States; Department of Chemical and Biomedical Engineering, FAMU-FSU College of Engineering, Florida State University, Tallahassee, FL 32310, United States; National High Magnetic Field Laboratory, Florida State University, Tallahassee, FL 32310, United States; Institute of Molecular Biophysics, Florida State University, Tallahassee, FL 32304, United States

## Abstract

Apolipoprotein A-I mimetic 4F, an 18-residue amphipathic α-helix, can self-assemble with lipids to form peptide nanodiscs, yet the molecular determinants governing their assembly and stability remain poorly understood. Here, using coarse-grained molecular dynamics (CG-MD), we capture the *de novo* formation of 4F nanodiscs with DMPC and reveal a multistep assembly pathway involving nucleation, fusion, and ellipse-to-disc maturation. All-atom back-mapping shows that the nanodisc rim is structurally heterogeneous and stabilized by aromatic-acyl interactions, Lys/Arg headgroup anchoring, and inter-peptide electrostatic contacts. Lipid composition and temperature critically regulate nanodisc integrity: DMPC supports continuous peptide belts and long-term stability, whereas DPPC below its main phase transition temperature suppresses fusion and yields fragmented, non-uniform rims. These findings validate the ability of CG-MD to resolve nanodisc assembly mechanisms. Experimental measurements corroborate the simulations, demonstrating that 4F nanodiscs exhibit lower thermal resilience than MSP nanodiscs while retaining structural integrity at moderate temperatures. As a functional benchmark, MSP nanodiscs inhibit amyloidogenic Aβ fibrillization, consistent with our earlier findings for 4F nanodiscs, indicating that a peptide-rimmed discoidal architecture is sufficient to suppress amyloid nucleation. Together, these results establish a mechanistic framework and design principles for single-helix peptide nanodiscs and delineate the conditions under which they converge with or diverge from MSP-based scaffolds.

## Introduction

Nanodiscs are disc-shaped lipid bilayers encircled by amphipathic scaffolds, forming a nanoscale membrane mimetic system in solution [1]. This distinctive architecture renders otherwise insoluble membrane proteins soluble in a native-like lipid environment, thereby preserving their structural integrity and biological function [1]. Nanodiscs overcome many limitations of traditional membrane mimetics [2,3] by eliminating detergents in the final reconstituted system [4] and minimizing curvature stress, thus providing a planar bilayer surface that closely mimics native biological membranes [5–7]. Owing to these advantages, nanodiscs have become a powerful platform for biochemical and structural studies of membrane proteins, enabling high-resolution techniques like cryo-EM and NMR for targets that previously required harsh detergent conditions. Beyond structural biology, nanodiscs are widely used to study membrane-associated processes, including the assembly of signaling complexes, and are increasingly explored in nanomedicine as delivery vehicles for drugs, lipids, or immunogenic complexes. The broad impact of nanodiscs in membrane science is evidenced by their widespread adoption and the emergence of numerous variants and alternative scaffolds built upon the same core design principle.

Compared to other membrane mimetics such as micelles, liposomes, and bicelles, nanodiscs offer several distinct advantages [8]. Detergent micelles often destabilize membrane proteins [4], whereas liposomes are inherently polydisperse and less suited for high-resolution structural methods. In contrast, nanodiscs provide a monodisperse, stable, and size-tunable lipid bilayer without requiring detergents [8–10]. Their lipid composition can be precisely controlled, and the flat discoidal geometry, which lacks the high positive curvature of small vesicles, better preserves native membrane protein conformations. Importantly, nanodiscs maintain a two-leaflet lipid bilayer structure, unlike micelles, enabling membrane proteins to be studied in a physiologically relevant lipid bilayer environment [11–17]. This native-like environment has proven essential for functional assays and drug discovery efforts. The success of the nanodisc concept has consequently inspired the development of numerous alternative membrane mimetic systems, including synthetic polymer- and peptide-based scaffolds [6,7,18,19], which aim to replicate the stability and versatility of the original protein-belt nanodiscs [8].

Apolipoprotein A-I (ApoA-I) provided the basis for the development of the first nanodisc scaffolds, given ApoA-I’s natural role in high-density lipoprotein (HDL) formation in blood. Truncations and mutants of ApoA-I, known as membrane scaffold proteins (MSPs), were engineered to self-assemble with lipids into nanodiscs of defined sizes, typically 8-16 nm in diameter. These MSP-based nanodiscs, which often contain two MSP molecules per disc, have become a useful tool in membrane protein research [14,15,20–23]. However, the expression and purification of relatively large MSPs can be resource-intensive, motivating the search for smaller, more tractable scaffold alternatives. One promising strategy has been the design of short amphipathic peptides that mimic the α-helical, lipid-binding regions of apolipoproteins. For example, the 18A peptide (18 residues long) is an early synthetic helix derived from the ApoA-I sequence and has been extensively studied as a potential HDL mimetic [24]. Multiple copies of such peptides can wrap around a lipid bilayer patch to form peptide nanodiscs. Indeed, recent studies have shown that tandem or multimeric versions of ApoA-I helices can form stable nanodiscs; for instance, linking two 18A helices yielded a bihelical peptide capable of assembling nanodiscs that incorporate membrane proteins in a detergent-free manner [25]. A notable development along these lines is the emergence of peptide nanodiscs, which employs multiple copies of a designed peptide to encapsulate membrane proteins without requiring large scaffold proteins [10]. In addition, nanodisc-forming peptides have been explored for isolating membrane proteins in nanodisc-like lipid assemblies [24,26,27]. Compared with traditional scaffold systems, peptide nanodiscs offer advantages in synthetic accessibility and design flexibility, while also reducing potential immunogenic concerns associated with large protein scaffolds.

Another approach involves the use of amphipathic polymers to generate nanodiscs. Styrene-maleic acid (SMA) copolymers, for example, can directly extract lipid patches from cell membranes to form polymer-encased lipid nanodiscs called styrene maleic acid lipid particles (SMALPs) [28–34]. In this process, the polymers insert into the membrane and excise small discoidal lipid-polymer assemblies. The SMA approach, along with newer SMA variants (including SMA-QA and SMA-EA), and styrene-free polymers (such as DIBMA and polymethacrylate derivative (PMA)) have gained prominence for detergent-free membrane protein extraction and purification. These systems yield stable, water-soluble lipid particles that retain native lipids and preserve protein function [31,34]. Each scaffold type presents distinct trade-offs. For example, MSP nanodiscs are well characterized and typically highly monodisperse, whereas peptide- and polymer-based nanodiscs are generally smaller and more synthetically accessible, enabling direct isolation from native membranes. Understanding how these different scaffolds stabilize lipid bilayers, and how they compare in terms of structure and stability, remains important for advancing nanodisc technologies.

Despite substantial progress in the experimental development of nanodiscs, the molecular mechanisms underlying their self-assembly remain difficult to resolve at atomic resolution. Experimentally, nanodiscs are typically formed by mixing scaffolds (proteins, peptides, or polymers) with lipids and sometimes in the presence of detergent, followed by detergent removal to trigger assembly. Capturing the stepwise assembly at atomic resolution is challenging because intermediate states are transient and heterogeneous, and commonly used techniques like cryo-EM or X-ray scattering generally report only ensemble-averaged end states. Experimental studies have shown that parameters such as temperature, scaffold-to-lipid ratio, and lipid phase state influence nanodisc formation pathways [35]. However, these approaches still lack a direct molecular view of how individual scaffold molecules and lipids coalesce into a discoidal bilayer. High-resolution techniques such as NMR and integrative structural modeling have further shown that nanodiscs are structurally heterogeneous than adopting a single static conformation. Complementary structural studies indicate that nanodiscs explore multiple conformational states and exhibit non-uniform lipid organization between the center and rim of the disc. This intrinsic dynamic complexity underscores the limitations of purely experimental approaches for tracking nanodisc assembly in real time and highlights the need for complementary molecular-level methods.

Computational approaches, particularly molecular dynamics (MD) simulations, have emerged as a powerful complementary tool for investigating the formation of nanodiscs [6,36–38]. MD simulations can provide a movie-like trajectory of nanodisc assembly and dynamics at molecular detail, which is exceedingly difficult to obtain experimentally. Early all-atom simulations of ApoA-I nanodiscs focused on equilibrium structure and dynamics, resolving how the scaffold protein belts arrange and how lipids diffuse within the disc [35,39]. For instance, all-atom MD runs revealed that ApoA-I’s helical belts can rearrange over hundreds of nanoseconds and that the disc can deform into elliptical shapes, an insight consistent with experimental observations [36]. However, such detailed simulations are computationally demanding and can suffer from kinetic trapping, it was noted that ApoA-I helices may require 10-20 µs to reach the correct arrangement on the disc, and shorter peptides ∼2-3 µs to migrate to the edge, meaning insufficient sampling or suboptimal initial conditions can lead to metastable, non-native states in simulations[40]. This highlights both the power and caveats of MD simulation, which can reveal slow collective motions and intermediate states; however, one must ensure adequate sampling and corroborate results with experiments. To accelerate the exploration of assembly pathways, coarse-grained (CG) MD simulations using simplified force fields, such as Martini, have been employed [5,41]. CG models sacrifice some atomic detail but drastically extend simulation timescales into tens of microseconds, where spontaneous self-assembly of nanodiscs can be observed [5,41].

We previously used CG-MD to capture the *de novo* formation of ∼7-8 nm diameter lipid nanodiscs within tens of microseconds. These simulations not only produced structural models consistent with experimental observations but also allowed us to probe properties such as nanodisc thermal stability through heating the systems. Notably, the MD-predicted optimal arrangement of scaffold molecules around the disc and the temperature at which discs disassemble correlated well with experimental data, providing strong evidence that the simulations capture realistic behavior. Simulations have further quantified how nanodisc size and geometric confinement influence lipid behavior. For example, lipid diffusivity and bilayer elasticity in nanodiscs differ from those in extended membranes, with nanodiscs generally exhibiting slightly stiffer bilayers due to their constrained geometry. These insights demonstrate the valuable synergy between simulation and experiment in nanodisc research, where MD can propose detailed molecular mechanisms and properties that can subsequently be verified and refined by experiments.

Amidst various nanodisc scaffold developments, ApoA-I mimetic peptides have garnered interest as minimalistic alternatives to full-length MSPs. One particular peptide, known as 4F, is a promising nanodisc-forming scaffold with added biomedical relevance. The 4F is an 18-residue amphipathic peptide (sequence: Ac-DWFKAFYDKVAEKFKEAF-NH_2_) with α-helical conformation originally designed to mimic the class A amphipathic helix of ApoA-I [42]. It contains four phenylalanine (F) residues on its hydrophobic face, hence “4F”, that enhance lipid-binding. This peptide was first studied for its atheroprotective effects [43], as it can bind lipids and improve HDL function; however, it has later been shown to self-assemble with phospholipids to form nanodiscs [44]. We demonstrated that 4F, in isolation or in a nanodisc state, exhibits anti-amyloidogenic activity against the Alzheimer’s amyloid-β (Aβ) peptide [6]. A mixture of 4F with different phospholipids forms nanodiscs smaller than 10 nm in diameter, which were able to significantly delay and suppress Aβ40 fibril formation but promote human amylin fibrillation [6]. Notably, 4F in its lipid-free state slowed Aβ aggregation, but the 4F-encased nanodiscs were even more effective, completely abolishing fibrillization at optimal peptide and lipid concentrations [45]. By comparison, conventional liposomes of similar lipid composition had substantially lower inhibitory efficacy on Aβ aggregation, underlining that the specific nanodisc format (with the peptide belt) is crucial for trapping Aβ or altering its aggregation pathway. These findings are important because they point to 4F-based nanodiscs that can have dual-function systems depending on the amyloidogenic protein they interact with, suggesting a therapeutic angle in diseases such as Alzheimer’s and type-2 diabetes.

Several other peptide scaffolds have been explored, but 4F offers a compelling combination of properties. For example, the 18A peptide and its derivatives form nanodiscs but often require chemical linkage of multiple helices or higher concentrations to achieve stable particles [10,27]. Chimeric peptides such as AEM28, which is a fusion of an ApoA-I helix with a segment that targets LDL receptors, have been shown to form nanodiscs [27]. The nanodisc scaffold peptide is another designed bihelical peptide that was successfully used to reconstitute membrane proteins [25]. While these newer peptides are promising, many are still being optimized for general use. The 4F peptide represents a compelling scaffold due to its small size, ease of chemical synthesis, and extensive prior pharmacological study. Unlike larger two-helical constructs, 4F is a single helix, meaning multiple copies are needed per nanodisc; however, this also offers flexibility, as adjusting the 4F:lipid ratio can tune the disc size [6]. Moreover, 4F’s proven ability to interact with membrane-active peptides such as Aβ adds a functional dimension that many other scaffolds lack. In terms of stability, initial reports indicate that 4F-based nanodiscs are quite stable in physiologically relevant conditions [27], although a detailed comparison with MSP nanodiscs has been lacking in the literature. Thus, 4F nanodiscs represent an attractive model system for investigating nanodisc assembly and stability, and for comparing peptide versus protein scaffolds head-to-head.

In this work, we combine CG-MD simulations with complementary experiments to elucidate the self-assembly of 4F nanodiscs and benchmark their properties against those of the well-established MSP nanodiscs. Using phosphatidylcholine (PC) lipids corresponding to DMPC or DLPC (referred to as DLPC/DMPC hereafter), which are represented identically in Martini CG models [46], together with the 4F peptide, we observe the spontaneous formation of nanodiscs *in silico*, enabling direct capture of the assembly pathway. We then compare the stability of 4F nanodiscs with that of MSP nanodiscs, including analyses of thermal melting behavior and size distributions measured by dynamic light scattering at various temperatures. These comparisons reveal how a small peptide scaffold withstands thermal stress and maintains disc integrity relative to a larger protein belt. Furthermore, by comparing 4F and MSP nanodiscs in terms of stability and interactions with Aβ, we identify key differences and advantages that not only validate the 4F nanodisc model but also inform the design of improved nanodisc platforms for biophysical and biomedical applications. Overall, this study demonstrates the capability of MD simulations to unravel the assembly of nanodiscs at the molecular level and underscores the potential of the apoA-I mimetic peptide 4F as a nanoscale membrane scaffold.

## Materials and methods

All experiments were performed in sodium phosphate buffer (pH 7.4) under the same sample-handling conditions as used in our previous studies [6]. Lipids were handled under inert conditions to prevent oxidation, and lipid stocks were prepared, quantified, and stored following previously established protocols [6].

### Coarse-grained molecular dynamics (CG-MD) simulations

CG-MD simulations were performed using the Martini force field. 4F peptides (modeled as amphipathic helices in Martini representation) were combined with PC lipids (DLPC/DMPC or DPPC) in explicit CG water with counterions to achieve overall electroneutrality. The initial configuration comprised randomized peptide and lipid positions (no pre-assembled disc geometry) to capture spontaneous self-assembly. Systems were energy-minimized and equilibrated under periodic boundary conditions. Production simulations were carried out at constant temperatures using a thermostat appropriate for Martini simulations and at constant pressure using semi-isotropic coupling to allow bilayer-like deformation. Self-assembly trajectories were run at 303 K to monitor formation and maturation, followed by extended simulations initiated from the final assembled nanodisc structure (snapshot at 11.4 µs) to assess stability at 310 K.

To probe thermal robustness, simulated annealing was performed by ramping the temperature from 310 K to 353 K over 6 µs, followed by a constant-temperature hold at 353 K for 14 µs while maintaining pressure coupling as described in our previous study [5]. Structural integrity was monitored using disc diameter, thickness, lipid loss events, and rim continuity metrics. Assembly intermediates and mature nanodiscs were characterized by (i) cluster analysis of peptide-lipid aggregates, (ii) radius/diameter and aspect ratio measurements, (iii) rim continuity (peptide coverage along the disc perimeter), (iv) lipid tail order proxies in CG coordinates, and (v) peptide tilt/azimuth distributions relative to the local disc plane. Contact analysis was performed using distance-based residue (bead) contact definitions to generate peptide-lipid and peptide-peptide interaction maps.

Representative CG snapshots (including the 11.4 µs assembled nanodisc) were back-mapped to an all-atom representation using standard Martini back-mapping workflows. Following back-mapping, atomistic structures were relaxed by restrained minimization and short equilibration to remove geometric artifacts prior to analysis. Peptide orientation distributions and lipid-contact enrichment were quantified from atomistic trajectories and structures using heavy-atom contact criteria for headgroup, glycerol, and acyl-chain regions and by computing helix tilt angles relative to the disc plane.

### Nanodisc preparation

4F peptide-lipid nanodiscs were assembled by spontaneous self-assembly under the same buffer and lipid handling conditions used previously [6,27]. DMPC lipids were combined with 4F peptide at the ratios specified in each experiment. Samples were equilibrated above the lipid phase transition as required for reproducible assembly, then clarified by low-speed centrifugation before downstream assays and size-exclusion chromatography (SEC) in working buffer.

MSP nanodiscs were prepared by standard cholate-assisted assembly followed by detergent removal, using MSP1D1-D73C expressed in *E. coli* and purified as described elsewhere [9], and DMPC under conditions consistent with established nanodisc protocols. After assembly, nanodiscs were purified by SEC and buffer-exchanged into the experimental buffer used across assays.

### NMR spectroscopy

4F and MSP nanodisc samples were prepared in 10 mM sodium phosphate buffer (pH 7.4) containing 5% D_2_O. Solution NMR spectra of nanodiscs were acquired on a Bruker 500 MHz NMR spectrometer equipped with a room-temperature TXI probe operating at 303 K (Billerica, MA, USA). Two-dimensional ^1^H-^1^H NOESY NMR spectra were recorded with mixing times (100 – 150 ms) to probe 4F- or MSP-lipid internuclear NOEs. Spectra were processed using standard apodization and baseline correction workflows in Bruker TopSpin software (4.5.0). NOE cross-peaks were assigned to lipid acyl-chain, headgroup, and peptide aromatic regions using chemical shift ranges and comparison across samples.

### Circular dichroism spectroscopy

Far-UV CD spectra were collected to quantify the α-helical content of 4F and MSP scaffolds in nanodiscs as a function of temperature. Wavelength scans were recorded under identical buffer conditions, and thermal ramps were monitored by ellipticity at 205 and 222 nm. Background spectra of buffer and lipid-only controls were subtracted. Temperature-dependent unfolding or dissociation signatures were inferred from loss of negative ellipticity at 208 and 222 nm and changes in spectral shape.

### Dynamic light scattering

Hydrodynamic size distributions of nanodiscs were measured by DLS using temperature-controlled measurements to compare thermal stability of 4F versus MSP nanodiscs. Samples were filtered immediately prior to measurement to remove dust. For thermal series, samples were equilibrated at each temperature before acquisition. Size distributions were reported as intensity-weighted profiles and tracked for shifts from monodisperse ∼10 nm nanodisc populations to larger assemblies (fusion or macrodisc formation) with increasing temperature, consistent with prior observations in peptide-based nanodiscs.

### Thioflavin-T (ThT) amyloid aggregation kinetics

Aβ40 aggregation kinetics were monitored using ThT fluorescence in the presence or absence of MSP DMPC nanodiscs across peptide:lipid ratios described in the Results. ThT was mixed with Aβ40 and nanodisc samples in the standard experimental buffer used previously. Reactions were plated in low-binding microplates in a BioTek plate reader, sealed to minimize evaporation, and incubated with controlled temperature and orbital shaking (as in prior reports [6]). Fluorescence readings were collected at regular intervals with excitation at 440 nm and emission at 485 nm, appropriate for ThT, and normalized to compare lag time, growth rate, and plateau intensity across conditions.

## Results and discussion

The spontaneous self-assembly of lipid-encircled nanodiscs is a complex, multi-stage process that unfolds on a microsecond timescale, making it particularly challenging to visualize at atomic resolution through traditional experimental techniques [47]. While methods like cryo-electron microscopy and X-ray crystallography provide exquisite detail of the final, stable particle, they are unable to capture th transient intermediates and the molecular rearrangements that occur during the formation pathway [48]. The CG-MD simulations presented here overcome this limitation, offering high-resolution and time-resolved insight into the complete self-assembly process of 4F nanodiscs.

Starting from a randomized mixture of 4F peptides and DMPC lipids at 303 K, the CG-MD trajectory captures a multistep self-assembly pathway that is broadly consistent with current understanding of nanodisc formation mechanisms from both experiment and simulation. Within the first microsecond, local peptide-lipid coalescence produces small mixed aggregates that resemble proto-discs or micelle-like clusters (**Figure 1**). These particles rapidly collide and coarsen, undergoing fusion events that yield elongated, ribbon- or ellipsoid-like particles within ∼7 µs. The elongated intermediates relax toward more regular, discoidal morphologies during 7-10 µs (**Figure 1**). The early coalescence and subsequent fusion steps mirror experimental observations that peptide nanodiscs can exchange lipids and merge in real time [49], indicative of mobile edges and low fusion barriers for peptide-stabilized rims.

**Figure 1.**
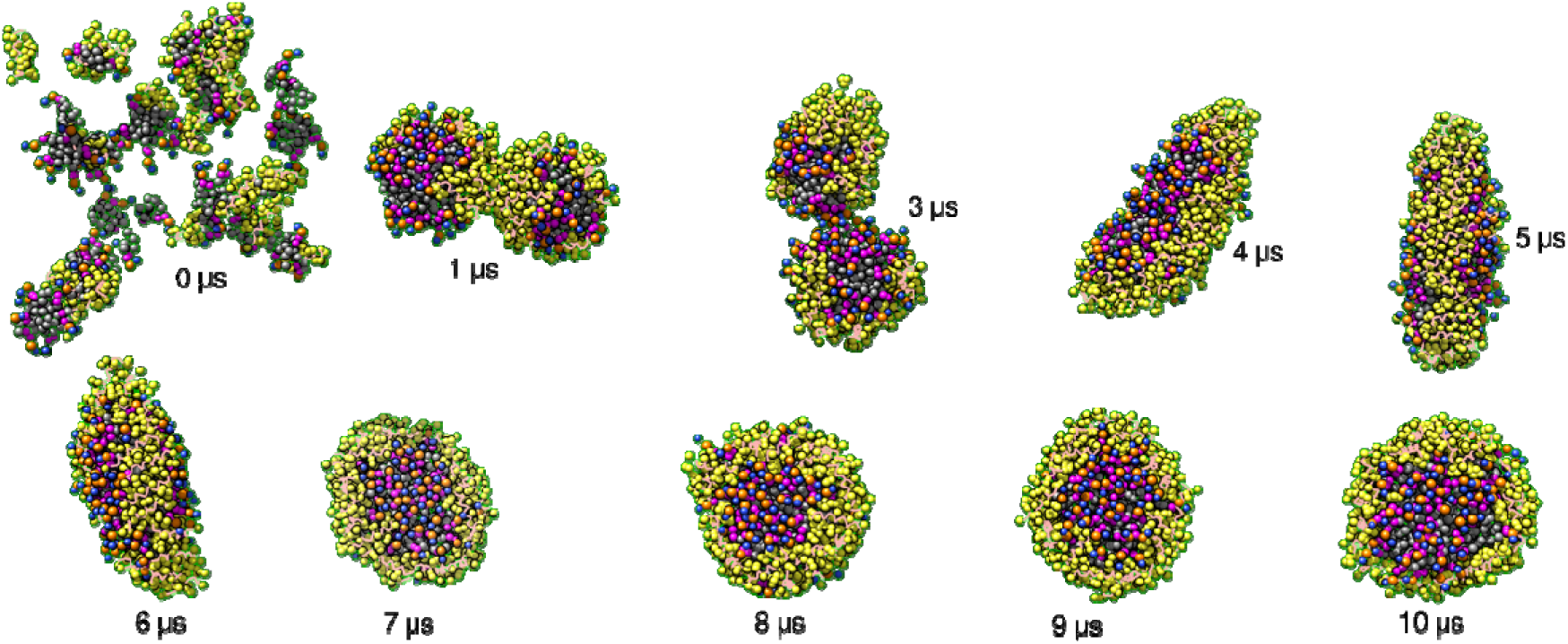
Dynamic self-assembly of 4F nanodiscs. CG-MD simulations capture the spontaneous formation of 4F nanodiscs over an 11.4-microsecond timescale. Snapshots, at key intermediate time points, illustrate the multi-step assembly mechanism. The initial state (0 µs) shows a random distribution of 4F peptides (yellow) and DMPC lipids (orange headgroups with purple acyl chains). Within the first microsecond (1 µs), small, irregularly shaped lipid-peptide aggregates form and begin to associate. These aggregates subsequently fuse and rearrange into an elongated, elliptical intermediate (3-7 µs) before undergoing a final morphological transition to a stable, circular nanodisc (8-10 µs). This simulation provides molecular-level insight into the kinetic pathway of peptide-nanodisc formation, a process challenging to resolve using conventional experimental methods.

Mechanistically, three features in the MD trajectory explain the ellipse-to-disc evolution. First, amphipathic 4F helices accumulate at the periphery, forming a quasi-continuous belt, which lowers the line tension of exposed bilayer edges and stabilizes the discoidal geometry. Second, lipids redistribute such that headgroups preferentially enrich at curved or edge regions while acyl chains align more orthogonally in the particle interior, an organization that has been reported for MSP nanodiscs and inferred from integrative SAXS/SANS/NMR/MD analyses showing elliptical fluctuations and radially varying lipid packing[35]. Third, inter-aggregate fusion shortens total rim length, trading anisotropic (elliptical) shapes for more circular discs to minimize edge energy, a trend also predicted by prior MD of nanodiscs and related nano-lipoprotein particles [36].

The timescales observed here (∼11 µs) are expected for Martini-level CG models and align with previous CG-MD studies in which spontaneous formation and maturation of nanodiscs or lipid-protein belts occurred within tens of microseconds, well beyond typical atomistic reach but fast enough to sample fusion or rearrangement events that are difficult to trap experimentally [50]. Importantly, the elliptical intermediates we observe (**Figure 1**) are not mere artifacts; small MSP nanodiscs are known to sample elliptical conformational ensembles at equilibrium, with aspect-ratio fluctuations captured by integrative scattering and MD [35]. Our peptide discs likely share the same elastic landscape, with progressive rim completion by 4F driving the ensemble toward rounder states.

These dynamics connect to functional behaviors reported for 4F systems. Prior work showed that 4F forms lipid nanodiscs and that such assemblies can fuse/exchange lipids experimentally, consistent with our coalescence pathway, and can modulate amyloid-β aggregation, implicating accessible peptide-lined rims and interfacial binding sites. The ease of rim remodeling seen here provides a physical basis for those observations, including mobile, peptide-rich edges that create binding surfaces and facilitate lipid/peptide exchange, features that differ from the more topologically constrained MSP belts. This mechanistic distinction complements recent reviews contrasting peptide, MSP, and polymer (SMA/DIBMA) discs, where scaffold chemistry governs edge elasticity, disc size distributions, and thermal robustness [51].

Coarse-graining accelerates kinetics and smooths energy barriers; thus, absolute times should be interpreted qualitatively. Likewise, we assumed identical CG bead mapping for DLPC/DMPC, while tail length differences minimally perturb Martini topologies, headgroup/tail parameterization nuances can shift preferred disc diameters and melting points. To address this point, we next performed CG-MD simulation using a similar setup for Martini model DPPC lipids. Furthermore, to assess the stability of the 4F-DMPC/DLPC nanodisc obtained at 11.4 µs on a longer timescale, we generated a simulation system that encapsulates the nanodisc structure (at 11.4 µs) as the initial structure and performed an additional 20 µs CG-MD simulation.

CG-MD simulations not only provide unprecedented insights into the assembly pathway of nanodiscs but also serve as a powerful tool to predict their stability and behavior under various physiological and thermal conditions. This is a critical advantage over experimental methods, which often provide only endpoint data on stability. The results presented in Figure 2 extend our understanding of the 4F-DMPC nanodisc system, demonstrating its remarkable long-term and thermal stability and revealing the critical role of lipid composition in its formation. Starting from the 11.4 µs endpoint in Figure 1, an additional 20 µs at 310 K shows a stable DMPC nanodisc with no rim fraying or lipid loss (**Figure 2A**). This behavior is expected for peptide-belted discs once a continuous amphipathic belt forms to lower edge line tension. This correlates to our previous study that shows variable-temperature DLS and ^31^P NMR on peptide nanodiscs can grow by fusion at elevated temperatures rather than disintegrate, consistent with a cohesive, mobile rim stabilized by many short helices [27].

**Figure 2.**
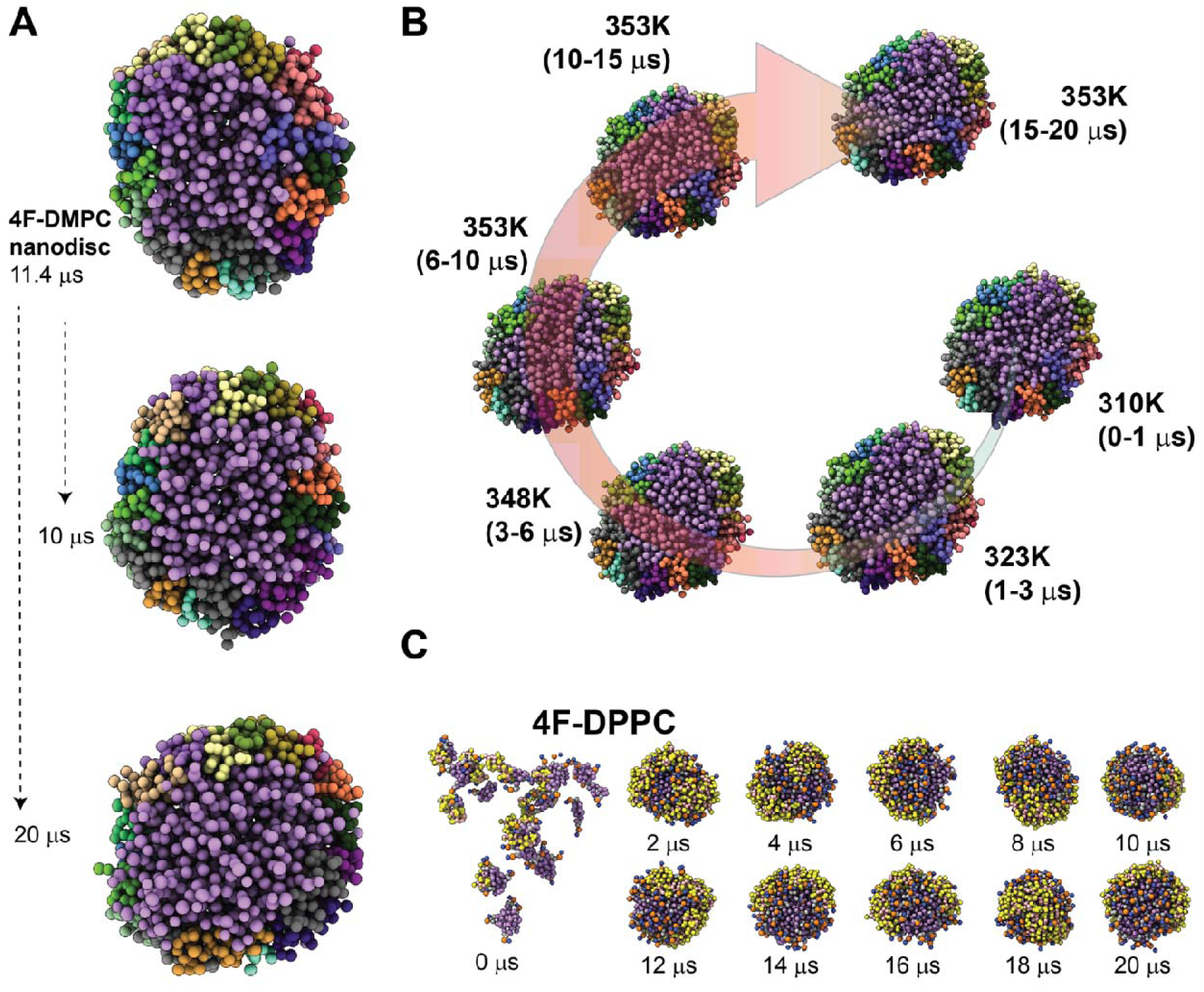
Stability and lipid-dependence of 4F nanodiscs in CG-MD. **(A)** Long-timescale stability of a 4F-DMPC nanodisc. The 11.4 µs endpoint from Figure 1 was used as the initial structure and simulated for an additional 20 µs at 310 K. The discoidal morphology and peptide belt remain intact, with no rim fraying or disintegration. This is consistent with peptide-nanodisc integrity and fusion behavior at elevated temperature *in vitro* [27]. **(B)** Thermal stability of 4F-DLPC/DMPC nanodisc probed using simulated annealing MD simulation. The 4F-DLPC/DMPC nanodisc was heated from 310 to 353 K over 0-6 µs and held at 353 K to 20 µs. The disc retains shape and rim continuity throughout. **(C)** Under identical conditions at 303 K, 4F-DPPC produces discoidal patches with non-uniform peptide coverage and no observable fusion-driven ellipse to disc maturation, consistent with DPPC being below its Tm (≈ 41–44 °C in vesicles/nanodiscs) and the known temperature requirement for efficient nanodisc assembly.

The thermal resilience of the 4F-DMPC nanodisc, a property not easily probed at high resolution experimentally, is further evidenced by the simulated annealing results (**Figure 2B**). Simulated annealing from 310 K to 353 K over 6 µs, followed by a 14 µs hold at 353 K. The nanodisc retains its shape and integrity even when heated to 353 K, well above the phase transition temperature (Tm) of DMPC lipid vesicles (23 °C or ∼296 K). This is consistent with the general observation that nanodisc lipids exhibit broadened, upward-shifted phase transitions relative to vesicles, yet the assemblies remain intact acros and above Tm. Our trajectory mirrors experimental reports of peptide-nanodisc fusion or size increase at higher temperatures without disintegration, reflecting lowered energetic penalties for peptide and lipid rearrangements as temperature rises [27,52].

When we replace DMPC/DLPC with DPPC, the system forms flatter discoidal or patch-like objects but lacks the robust rim uniformity, rim-peptide continuity, and the ellipse-to-disc maturation seen in Figure 1. At 303 K, DPPC is below its gel to fluid transition (Tm ≈ 41 °C in vesicles and ≈ 44 °C in nanodiscs), so chains remain ordered and lateral mobility is reduced, which suppresses protodisc fusion and hinders rim annealing into a uniform belt (**Figure 2C**). This observation fits together with experimental guidance that nanodiscs assemble most efficiently near/above the lipid Tm. Together, these data explain why 4F readily forms well-rimmed discs with DLPC/DMPC but yields fewer uniform peripheries and fewer fusion derived elliptical intermediates with DPPC under the same conditions. We explain that for shorter-chain PCs (DLPC/DMPC), the fluid state at 303 K supports rapid peptide diffusion and edge-line-tension minimization, enabling the ellipse to disc pathway and long-time stability. For DPPC, the gel-like state at 303 K stiffens the bilayer and frustrates rim remodeling and therefore only discoidal patches emerge, with non-uniform peptide coverage. These trends are fully consistent with (i) the identical Martini mapping of DLPC and DMPC versus the distinct, longer DPPC mapping, and (ii) the known Tm shifts and cooperativity changes when lipids are confined in nanodiscs [53].

The formation of a stable, disc-like structure in the 4F-DMPC nanodisc is driven by a combination of a hydrophobic core and a polar "belt" of charged residues, a mechanism well-documented for amphipathic peptides that mimic apolipoprotein A-I. Back-mapping the 11.4 µs CG snapshot to all-atom and quantifying contacts reveals that 4F helices around the DMPC/DLPC nanodisc rim do not adopt a single angle (**Figure 3A**). Instead, they populate a broad distribution of tilt/azimuthal orientations relative to the local disc plane, where some lie nearly parallel, and others are more oblique. This heterogeneity is expected for peptide-belt nanodiscs and reconciles well with prior structural work. Oriented ssNMR on ApoA-I mimetic helices (14A/18A) showed shallow, in-plane-biased alignment with small average tilt angles when discs form, but those measurements report ensemble averages, not a single rigid geometry [54]. SAXS/SANS with modeling on 18A-DMPC discs likewise recovered discoidal particles with peptide belts comparable to ApoA-I/MSP discs, while allowing shape and orientational flexibility of the belts [55]. Moreover, integrative SAXS/SANS/NMR/MD analyses of MSP nanodiscs established that even “canonical” belts exhibit elliptical fluctuations and heterogeneous lipid organization, implying that belt segments explore a range of orientations on microsecond timescales [35]. That ensemble view aligns with the mixed orientations observed in our simulation (**Figure 3A**).

**Figure 3.**
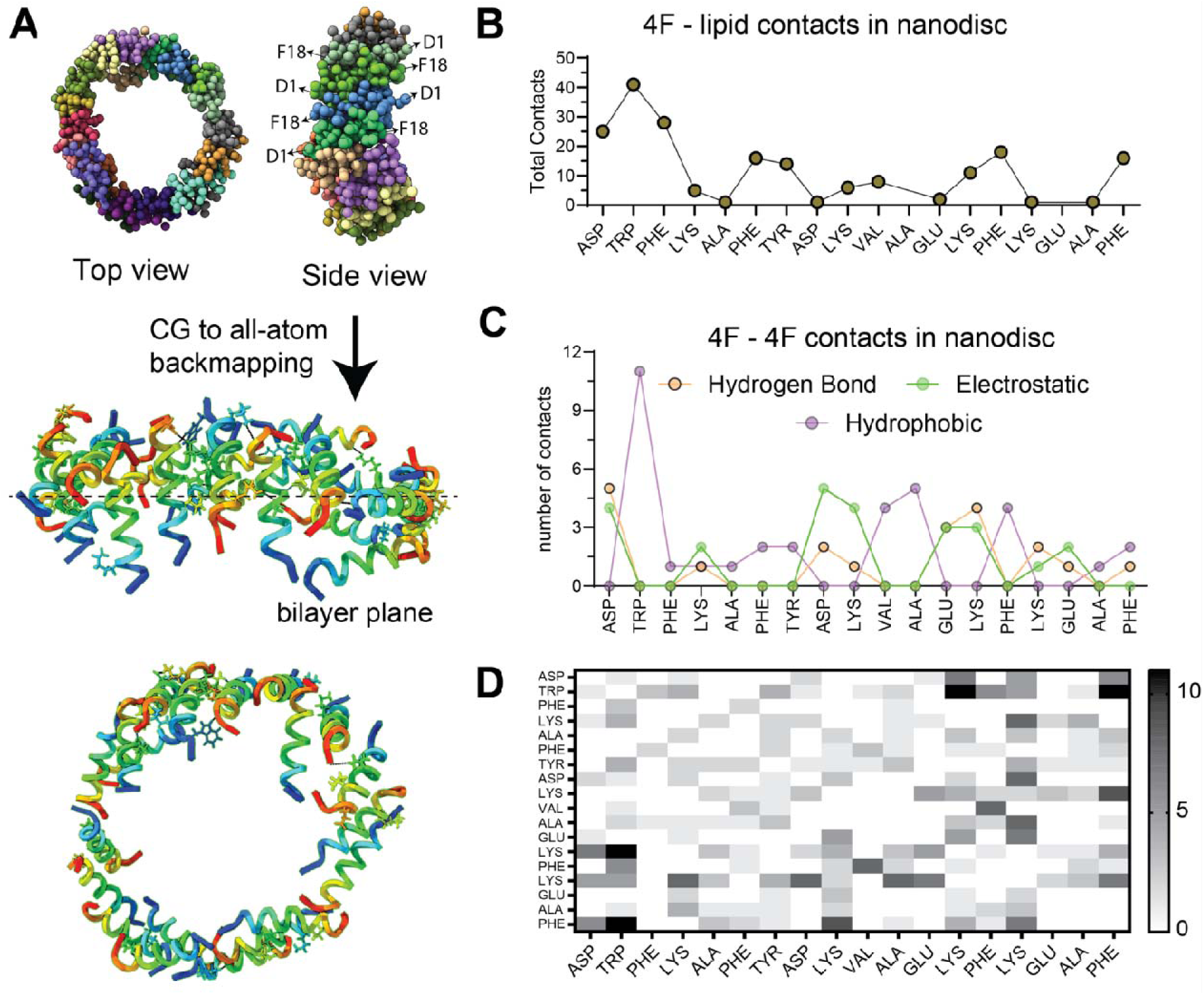
All-atom contacts explain a distribution of 4F orientations in the nanodisc. **(A)** CG to all-atom back-mapped snapshot of a 4F-DMPC/DLPC disc shown as CG-bead (top) and ribbons (side/top view) on bottom. Helices around the rim exhibit mixed orientations where some are nearly parallel, other more tilted forming a flexible belt. The dashed line in the all-atom structure represents bilayer plane axi and intermolecular peptide residues involved in salt-bridge and hydrogen bonding are shown in sticks. **(B)** Per-residue peptide-lipid contact counts highlight strong hydrophobic (Phe/Leu/Trp-acyl) and electrostatic (Lys/Arg-phosphate/glycerol) interactions that anchor helices at the interface. **(C)** Peptide-peptid contacts involving hydrogen bonds, hydrophobic interactions, and electrostatic interactions reveal head-to-tail associations between neighbouring helices, consistent with the antiparallel pairing observed for ApoA-I mimetics. **(D)** Residue-residue contact heatmap between peptides emphasizes terminal and mid-face contact clusters. Together, these contacts stabilize the rim while allowing orientational heterogeneity, in line with experimental and MD studies of peptide/MSP nanodiscs [54,55].

Our contact analysis further led us to rationalize why multiple orientations are simultaneously stable (**Figure 3**). First, 4F’s hydrophobic face (Phe/Leu/Trp) shows frequent contact with acyl chains, anchoring the helix into the rim (**Supplementary Figure S1**). This explains how 4F differs subtly from it 2F progenitor by deeper aromatic engagement and additional headgroup contacts to form nanodiscs, consistent with solution NMR observations on 4F:DMPC discs [56]. Second, basic residues (Lys/Arg) repeatedly contact phosphate/glycerol groups, pinning helices near the interface. The balance between tail and headgroup interactions permits multiple local minima, generating parallel versus more tilted poses depending on local curvature and lipid packing[56]. Third, the residue-residue map shows inter-peptide links concentrated near opposite termini, consistent with head-to-tail antiparallel associations reported for ApoA-I mimetics (22A) and inferred for peptide discs (**Figure 3A**). The electrostatic pairs, for example, Lys-Asp/Glu and lateral hydrophobics, stabilize neighboring belt segments without enforcing a single strict geometry [57].

The contact analysis of the 4F peptides further revealed a cooperative stabilization of the nanodisc rim driven by both polar and nonpolar interactions (**Figure 3B**). Binary contact maps show that positively charged residues (Lys4, Lys13, Lys15) make the highest number of unique inter-peptide contacts, frequently partnering with Asp/Glu to form hydrogen bonds and salt-bridge networks that “stitch” adjacent peptides. Aromatics with bulky hydrophobic sidechains (Trp2, Phe18, Tyr7) contribute numerous nonpolar contacts, consistent with π–π/CH–π packing at the solvent-exposed rim. The coexistence of electrostatic steering and hydrophobic anchoring provides a mechanistic basis for the spontaneous discoidal assembly observed in CG simulations, and mirrors prior findings for amphipathic peptide scaffolds that mimic MSP behavior while offering tunable residue-level control over stability and size. Such insights suggest that residue-level engineering, particularly of charged and aromatic hotspots, can be exploited to design nanodiscs with enhanced stability and tailored membrane-protein compatibility.

Our coarse-grained MD simulations (**Figure 3**) revealed that MSP and short-peptide scaffolds stabilize nanodiscs through distinct mechanisms. The MSP protein forms a continuous double-belt around the lipid patch [57], whereas multiple 4F peptides assemble a more flexible, discontinuous rim. In simulations, the short amphipathic helices of 4F exhibited heterogeneous tilt orientations and fewer inter-helix contacts, suggesting a less cooperative hold on the disc edge. In contrast, the two covalently linked helices of MSP (mimicking ApoA-I’s paired helices) provided a more uniform, constrained belt, a feature associated with greater nanodisc stability [58]. Indeed, prior studies show that peptide-only nanodiscs tend to be less stable, for example, the 18A peptide-DMPC nanodiscs gradually reorganize into larger particles over time [57] and lose structure above the lipid’s Tm as peptides migrate off the rim [58]. By tethering helices together through Pro linkers or disulfide bonds, researchers have improved peptide-disc stability, underscoring that a laterally associated “double belt” architecture is crucial for maintaining disc integrity [58]. Consistent with this, rigid dimeric peptides (with no or short linkers) form more structurally stable nanodiscs than highly flexible variants [57]. These insights led us to predict that MSP-based nanodiscs would withstand thermal stress better than 4F-based nanodiscs, which lack a contiguous belt and exhibit higher intrinsic hydrophobicity.

To experimentally evaluate the lipid contacts and orientational heterogeneity predicted by the MD simulations, we performed 2D ^1^H-^1^H NOESY NMR measurements for 4F:DMPC nanodiscs and compared them with MSP:DMPC nanodiscs (**Figure 4**). The 2D NOESY spectrum and its corresponding projections display a combination of narrow and broad lines for 4F nanodiscs. Strong NOE signals were observed for the DMPC acyl chain protons (∼0.5-1.6 ppm) and the glycerol/headgroup protons (∼3.0-4.5 ppm), as well as the 4F aromatic protons (6.6-7.3 ppm), while weak signals were observed for the 4F backbone amide protons (7.4-8.8 ppm) (see 1D projections in **Figure 4A**). Consistent with the MD-derived contact maps, the 4F nanodiscs displayed a broader distribution of peptide-lipid NOE cross-peaks, particularly between 4F aromatic protons and DMPC headgroup and upper-chain methylene groups, indicating efficient NOE transfer due to strong lipid-peptide interactions. In contrast, cross-peaks involving non-aromatic protons were significantly weaker. These findings support the CG MD results, demonstrating that the aromatic residues of the 4F peptide predominantly interact with lipids within the nanodiscs. The NOEs correspond to the same phosphate or glycerol and acyl-chain contacts enriched in the simulation-derived per-residue interaction profiles, supporting an interfacial helix orientation in which 4F samples both shallow (near-parallel) and more tilted poses relative to the bilayer plane. Furthermore, the Trp2 indole NOE cross-peak with DMPC acyl chain is consistent with its predicted anchoring role in the MD back-mapped structures (**Figure 4A and Supplementary Figure S2**). In contrast, MSP nanodiscs exhibit fewer peptide-lipid NOEs (**Figure 4B**), consistent with their more continuous and structurally constrained double-belt architecture, which restricts local helix mobility. Thus, the NOESY patterns provide experimental support for the MD-derived model of a flexible, heterogeneously oriented 4F rim stabilized by mixed hydrophobic and electrostatic contacts and highlight a fundamental structural distinction between peptide- and protein-based nanodiscs.

**Figure 4.**
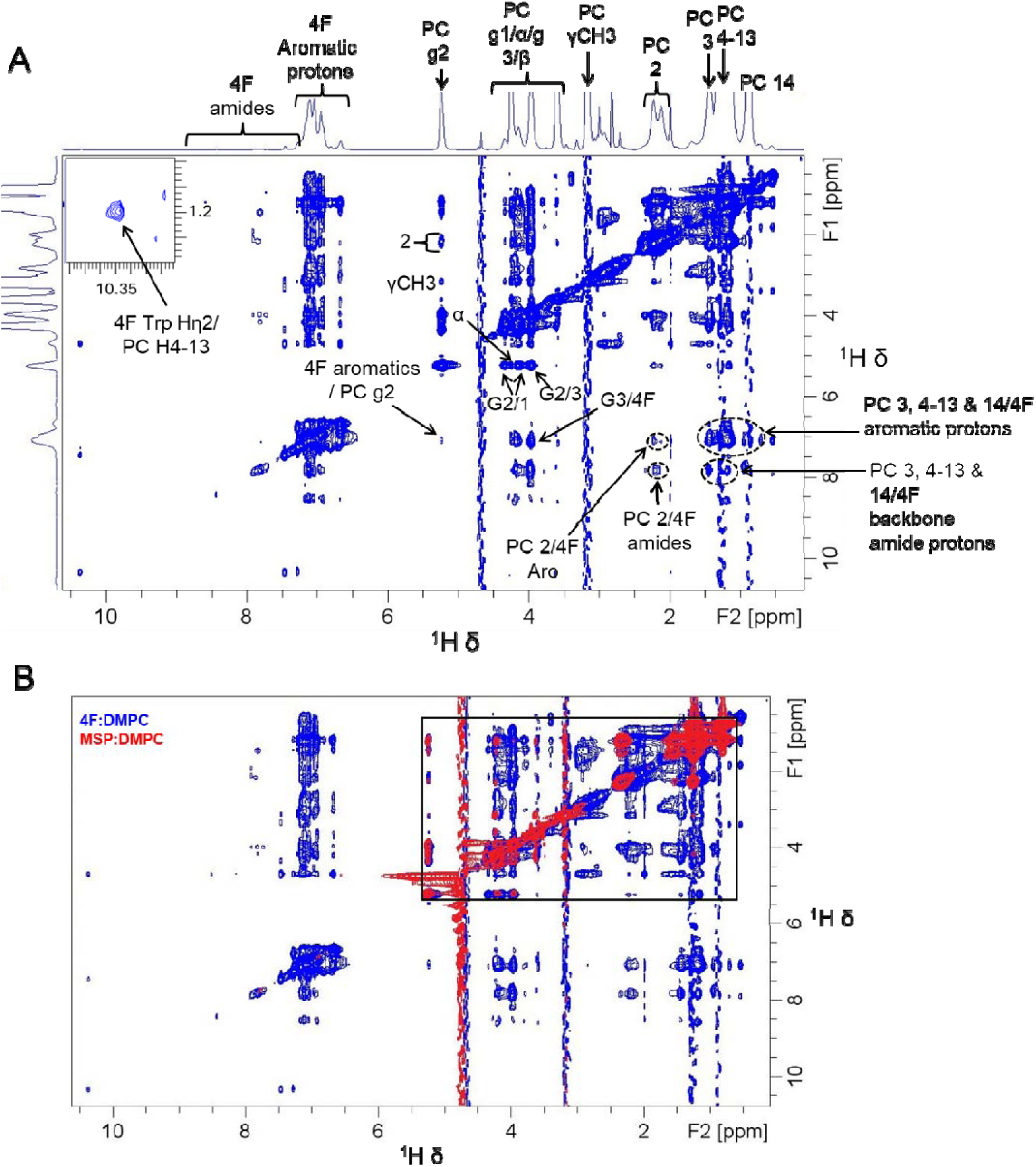
2D ^1^H-^1^H NOESY spectra of 4F:DMPC and MSP:DMPC nanodiscs. (**A**) 2D NOESY NMR spectrum of the 4F-DMPC (5 mg/mL) nanodiscs (blue) showing resolved amide, aromatic, and aliphati resonances from 4F together with phosphocholine headgroup and acyl-chain signals of DMPC. Cross-peaks between 4F and DMPC protons are observed in regions corresponding to lipid headgroup (g2, g1/α/β) and upper-chain methylene groups. (**C**) The NOE cross-peak between downfield 4F Trp indol amide (∼10.35-10.40 ppm) and DMPC acyl chain (1.1 - 1.3 ppm), which reports on local contacts within the nanodisc environment, is expanded. (**B**) Overlay of 2D NOESY NMR spectra of 4F:DMPC (blue) and MSP:DMPC (red) nanodiscs (5 mg/mL) highlights that the 4F nanodiscs exhibit a higher number of observable NOE cross-peaks than MSP nanodiscs (boxed). The NOEs in the aromatic/amide region are observed only for the 4F nanodiscs. NMR spectra were recorded on a Bruker 500 MHz spectrometer equipped with a TCI probe operating at 303 K (above the T_m_ of DMPC).

Experimental design puts these predictions to the test by comparing the thermal behavior of MSP1D1-D73C with 4F nanodiscs using DMPC lipids by dynamic light scattering (DLS) and circular dichroism (CD) spectroscopy (**Figure 5**). DLS tracks the particle size distribution as temperature rises, while CD monitors the α-helical content of the scaffolds, thus together probing the discs’ structural integrity under thermal stress. Literature shows that intact nanodiscs maintain a monodisperse ∼10 nm diameter by DLS, but upon heating, they can fuse into larger “macrodiscs” or aggregates, causing a growing hydrodynamic radius and multi-peak size distributions [27]. Simultaneously, the scaffold’s helices may unfold or dissociate, seen as a drop in CD ellipticity at 222 nm (loss of α-helix). For example, peptide nanodiscs typically begin enlarging and losing stability around 55-65 °C [27], whereas MSP-based nanodiscs often remain intact until higher temperatures, as the MSP belt itself only unfolds near ∼70-80 °C in some cases [59]. We therefore tested our MD-derived hypotheses. If 4F nanodiscs have a lower thermal limit, we expect their DLS profile to broaden or shift to larger sizes at higher temperatures than that of MSP-D73C discs, and their CD signal to denote earlier helix disruption. A sharper thermal transition for 4F would indicate a more cooperative mechanism, whereas MSP’s extended stability would reflect its robust, covalently constrained belt [59]. Our experimental data confirms this stabilizing role of scaffold architecture. By demonstrating how MSP-D73C retains a small, monodisperse size and helical structure up to higher temperatures than the 4F system (**Figure 5**), our data support a model of cooperative rim stabilization and validate the simulation insights from Figures 1-3.

**Figure 5.**
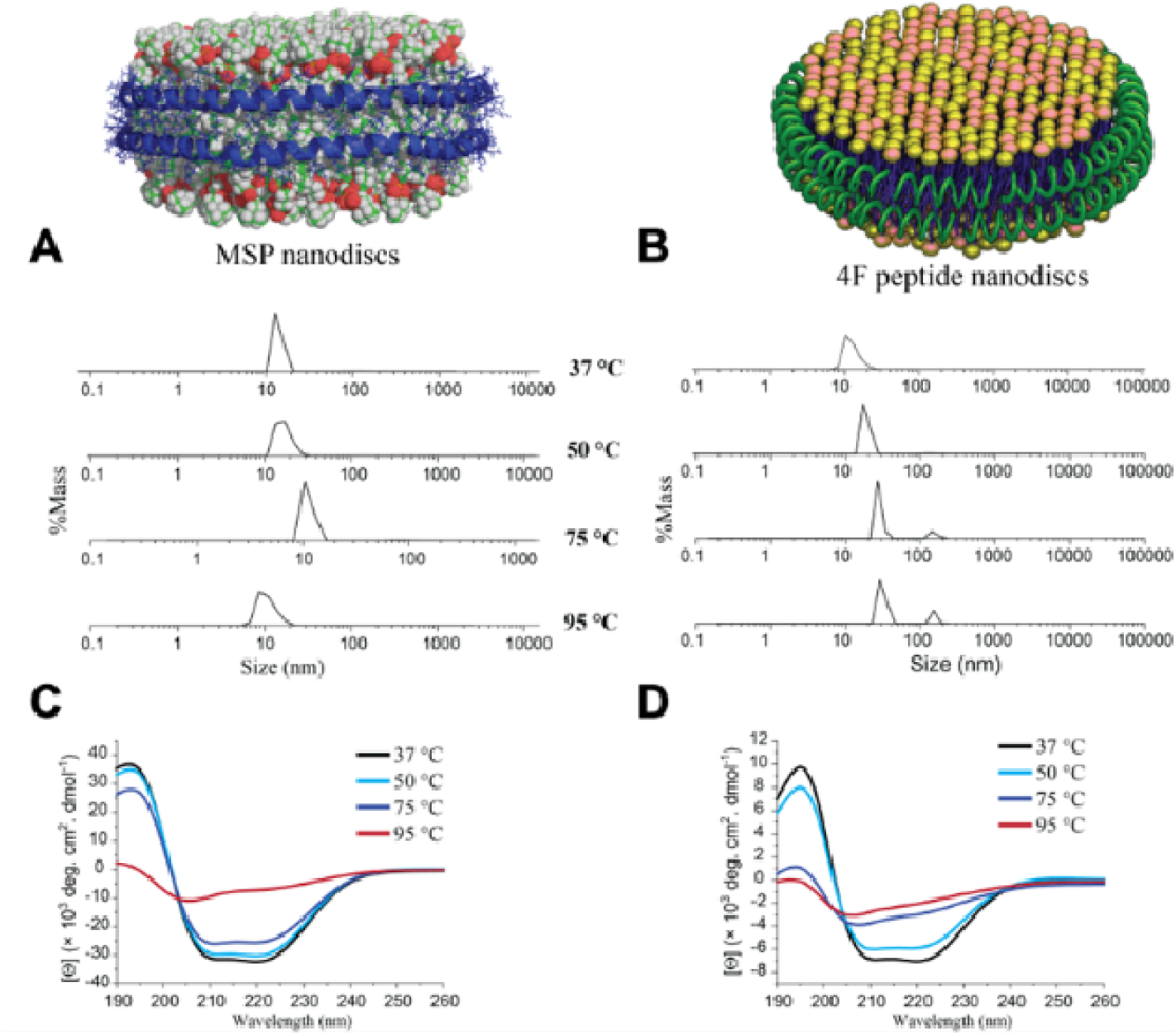
Thermal response of MSP and 4F peptide DMPC nanodiscs. **(A-B)** Dynamic light scattering (DLS) size distributions (hydrodynamic diameter) for MSP1D1-D73C DMPC **(A)** and 4F-DMPC **(B)** nanodiscs recorded at 37, 50, 75, and 95 °C. MSP nanodiscs remain narrowly distributed and near their initial size at 37-50 °C, with only modest broadening by 75 °C, and extensive growth/polydispersity emerges primarily at 95 °C. In contrast, 4F nanodiscs exhibit earlier temperature sensitivity, with distributions broadened by 50 °C and a shift to larger apparent sizes by 75 °C, consistent with heat-induced fusion/aggregation, and multiple populations dominate at 95 °C. **(C-D)** Far-UV circular dichroism (CD) spectra of MSP1D1-D73C/DMPC **(C)** and 4F/DMPC **(D)** nanodiscs at the same temperatures. Both scaffolds are α-helical at 37 °C (minima near 208/222 nm). Upon heating, MSP retains substantial ellipticity through 75 °C and shows partial loss by 95 °C, whereas 4F displays an earlier reduction of the 222 nm band (by 50-75 °C), approaching near random coil at 95 °C.

To test whether the functional readout of an amphipathic belt is conserved irrespective of scaffold architecture, we asked if MSP nanodiscs also suppress Aβ aggregation, a property we previously established for 4F nanodiscs [6]. Normalized ThT kinetics show that MSP-DMPC nanodiscs strongly delay and attenuate Aβ_1-40_ fibrillization across peptide:lipid ratios (1:20 to 1:100), whereas Aβ alone rapidly reaches a high plateau (**Figure 6**). Despite the distinct rim organizations revealed in Figures 1-3 (multi-helix, orientation-heterogeneous 4F versus contiguous MSP belt), both scaffolds present lipid-peptide edges that sequester early Aβ species and divert nucleation. Thus, the aggregation study functionally validates the central mechanistic insight from our simulations, which is that a discoidal bilayer with an amphipathic rim, even when built from a single-helix peptide, is sufficient to inhibit Aβ self-assembly.

**Figure 6.**
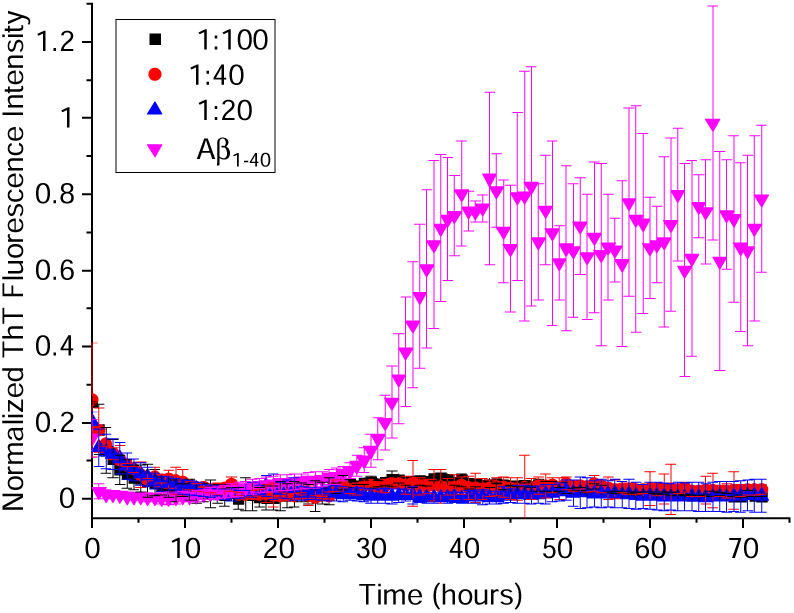
MSP nanodiscs inhibit Aβ_1-40_ fibrillization. ThT (thioflavin T) fluorescence kinetics of Aβ_1-40_(control, magenta) and Aβ co-incubated with MSP-DMPC nanodiscs prepared at the indicated Aβ:lipid stoichiometries. Curves are normalized, and data points show mean ± SD over replicates. MSP nanodiscs extend the lag phase and reduce the plateau signal relative to Aβ alone, indicating suppressed fibril formation. This correlates with our prior result that 4F nanodiscs inhibit Aβ aggregation [6], and support a shared interfacial sequestration mechanism.

## Conclusion

This work establishes that a minimal, single-helix ApoA-I mimetic (4F) peptide can spontaneously assemble with PC lipids into stable nanodiscs. CG-MD simulations directly capture this process as a multi-step pathway involving nucleation, fusion, and ellipse-to-disc maturation. We further identify key lipid and temperature dependencies: DMPC and DLPC lipids support uniform rim formation and long-time structural integrity, whereas DPPC below its main phase transition temperature (T_m_) impedes fusion and yields non-uniform rims. These findings validate CG-MD as a reliable tool for probing nanodisc assembly *in silico*. Despite architectural differences, both ApoA-I mimetic peptide 4F and the canonical membrane scaffold protein (MSP1D1) can be engineered into stable lipid nanodiscs that potently inhibit Aβ aggregation [6,60]. This observation suggests a common mechanism whereby amphipathic helical belts bind and stabilize early Aβ species, preventing their progression into pathogenic fibrils, consistent with our previous hypothesis [6]. Our results therefore highlight that amphipathic belt proteins from diverse origins can be tailored as nanodisc-based inhibitors of amyloid formation. Altogether, this study provides mechanistic insights for designing single-helix peptide nanodiscs and defines their functional operating envelope relative to MSP scaffolds.

## Acknowledgement

This study was supported by NIH (DK132214 to A.R.) and NSF (DMR-2128556).

