## Supporting Information for "Molecular Mechanisms Governing Peptide Nanodisc Assembly and Stability"

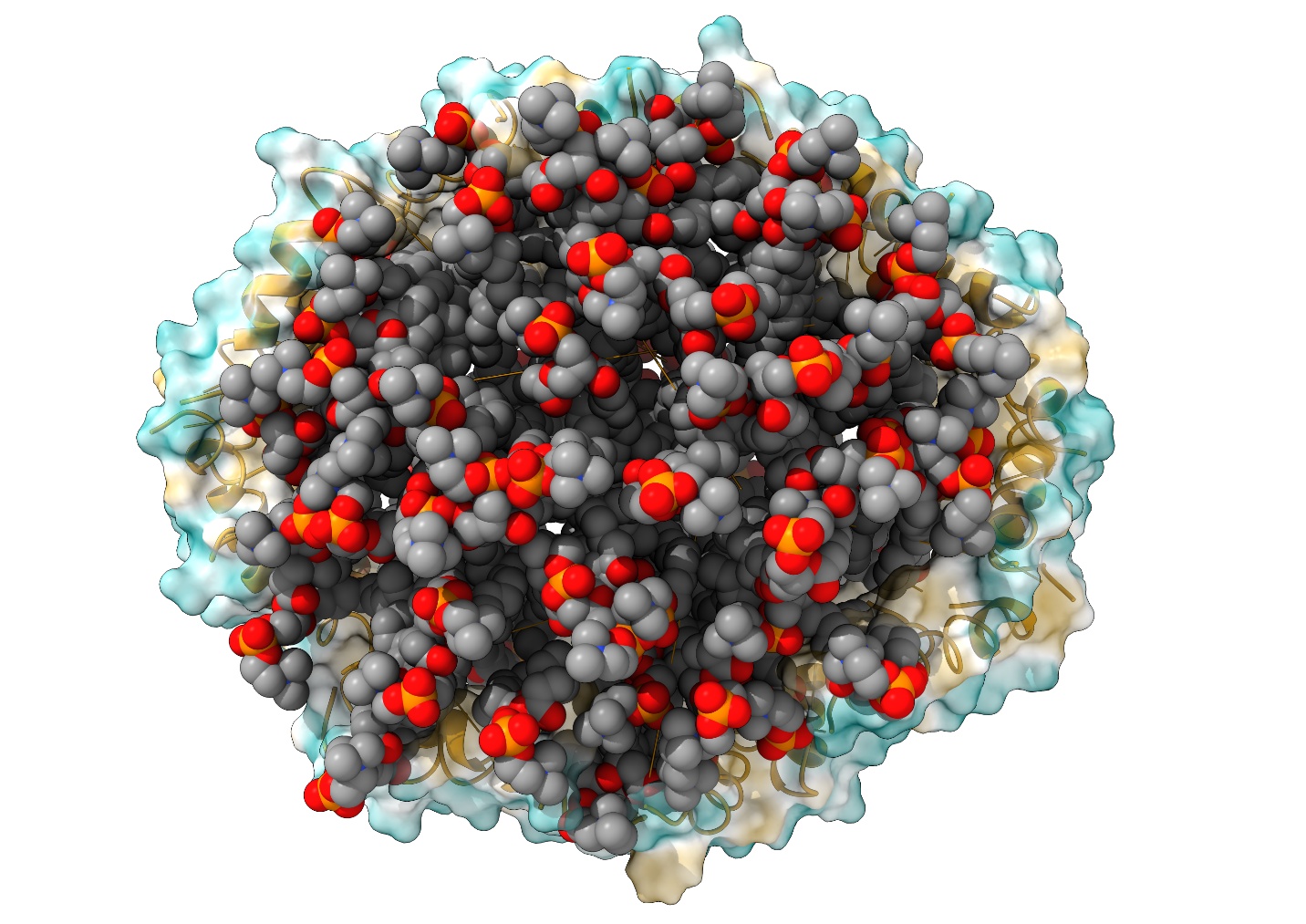

**Figure S1**. Final molecular dynamics snapshot of the self-assembled 4F peptide nanodisc. Peptides are shown as cartoon and surface representations forming a continuous rim that encapsulates the DMPC lipid core (spheres). Hydrophobicity coloring (orange: hydrophobic; blue: hydrophilic) highlights segregation of lipid-facing hydrophobic residues and solvent-exposed polar residues.

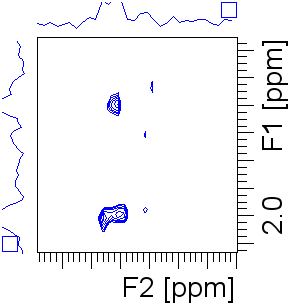

1.2

2.0

10.4

10.2

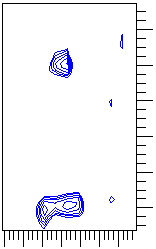

10.36

2.0

1.2

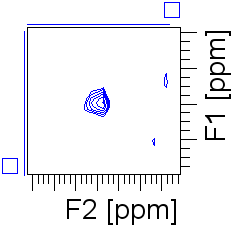

10.35

1.2

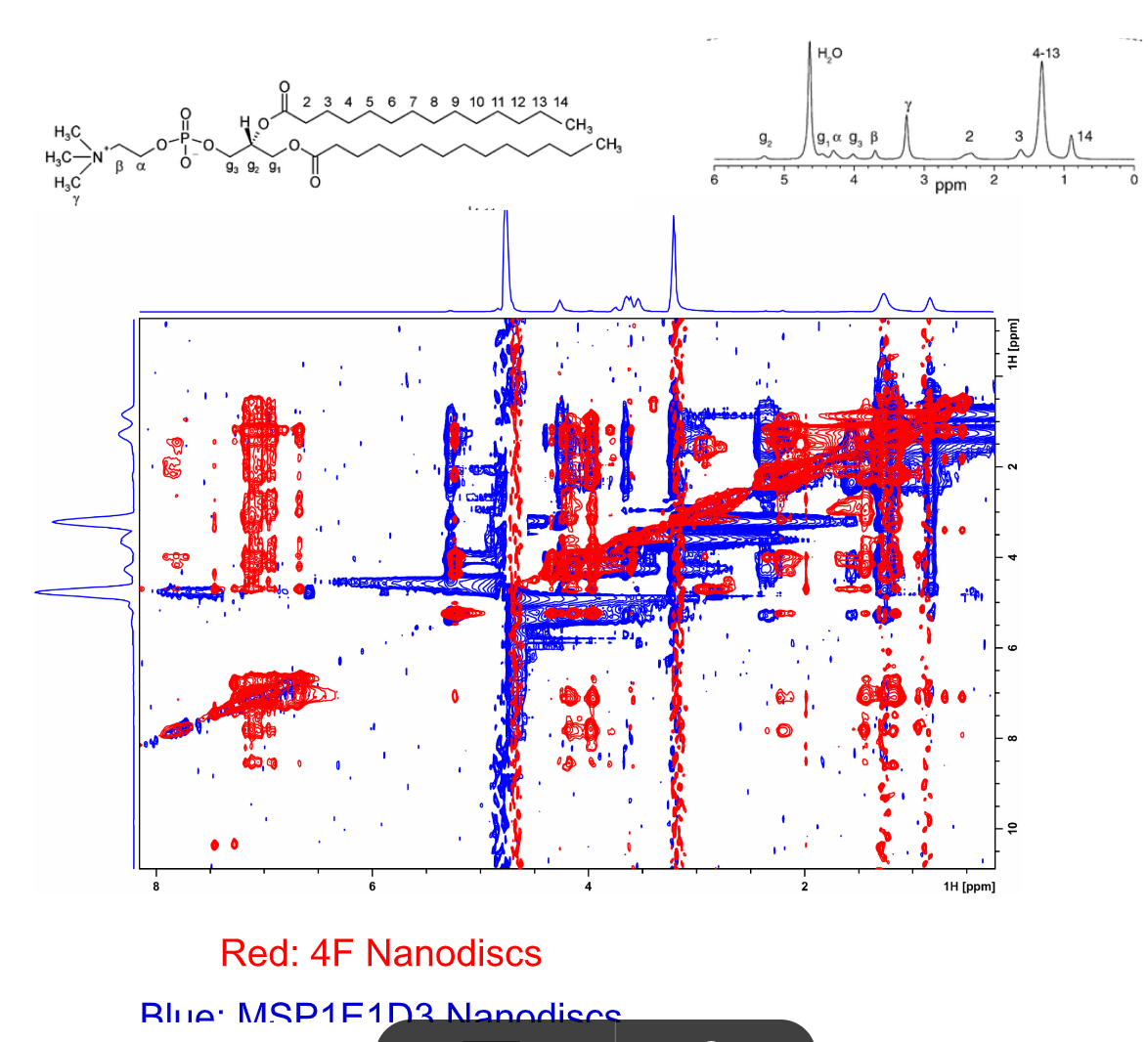

1H (ppm)

1H (ppm)

1H (ppm)

1H (ppm)

1H (ppm)

1H (ppm)

**Figure S2**. Assignment and NOE validation of 4F peptide-DMPC interactions. The top panel shows the chemical structure of DMPC and its corresponding 1D ^1^H NMR spectrum with labeled proton resonances. The bottom panels present expanded regions of the 2D NOESY spectrum highlighting intermolecular NOEs between the indole NH proton of the 4F Trp residue (~10.3 ppm) and DMPC lipid acyl chain methyl protons (~1.2-1.4 ppm). These cross-peaks indicate close spatial proximity (<5 Å) between the aromatic residue and lipid chains, supporting insertion of the peptide into the lipid environment.
